# Whole mitochondrial genome analysis of *Aedes aegypti* reveal association with *Wolbachia* infection

**DOI:** 10.1101/2024.12.04.626768

**Authors:** Bhavna Gupta, Melveettil Kishor Sumitha, G Navaneetha Pandiyan, Mariapillai Kalimuthu, Rajaiah Paramasivan, Manju Rahi

**Affiliations:** ICMR-Vector Control Research Centre (VCRC), Field Station, Madurai, Tamil Nadu, India, 625002; ICMR-Vector Control Research Centre (VCRC), Puducherry, India, 605006

**Keywords:** *Aedes aegypti*, mitochondrial genome, *Wolbachia*, India, maternal inheritance, mitochondrial lineage

## Abstract

The mitochondrial genomes (mitogenomes) of nine *Aedes aegypti* samples from India were analysed along with 34 mitogenomes from global samples retrieved from GenBank. The mitogenome size of Indian samples ranged from 15,730 bp to 16,374 bp. A total of 199 genetic variants were identified among Indian samples, with the majority (90%) occurring in protein-coding genes, followed by rRNA and tRNA genes. Phylogenetic analysis of the 43 genomes revealed two major clades. The similar clustering pattern was observed in the traditional mitochondrial markers for which extensive global data is available, indicating that individual mitochondrial markers of *Ae. aegypti* share the common genealogy as reflected by the complete mitogenome. In addition to exploring genetic diversity, we investigated the relationship of these two mitochondrial clades with *Wolbachia* infection. Our analysis revealed that *Wolbachia*-infected samples were predominantly located within one of the mitochondrial clades, suggesting a potential association between specific mitochondrial lineage and *Wolbachia* infection. This analysis demonstrates the extent of genetic diversity in *Ae. aegypti* mitogenome and highlights how this diversity is associated with *Wolbachia* infection, a maternally inherited endosymbiont. These findings have implications for the effectiveness of *Wolbachia*-based mosquito control strategies.

## Introduction

Mitochondrial markers are widely used for species identification, molecular phylogenetics, and evolutionary studies. This popularity has arisen due to several key characteristics of the mitochondrial genome, including its high substitution rates, uniparental inheritance, minimal recombination, and the presence of multiple genome copies (Desalle et al., 2017; Galtier et al., 2009). Several mitochondrial genes such as cytochrome c oxidase subunit I (COI), Nicotinamide-adenine dinucleotide (NADH) dehydrogenase subunit 2, 4, 5 (ND2, ND4, ND5), and cytochrome b (Cyt b) have been used extensively for such studies (Bušić et al., 2024; Chan et al., 2014; Kumar et al., 2022; McFadden et al., 2004; Wang et al., 2021; Yu and Zhang, 2021). These genes are known to share a common genealogy due to their linkage within the mitochondrial genome. Moreover, the proteins encoded by mitochondrial genes play essential roles in oxidative phosphorylation and energy metabolism, they are more susceptible to non-neutral evolutionary pressures. However, mitochondrial genes have been observed to exhibit distinct phylogenetic signals because of the adaptive selection pressures acting upon them (Ramos et al., 2020; Zhang et al., 2021). Thus, combining multiple genes or analyzing the entire mitochondrial genome are generally preferred. This can provide detailed insights on evolutionary history as well as adaptive evolution across species.

*Aedes aegypti*, a primary vector for several arboviral diseases including dengue, Zika, chikungunya, and yellow fever, is a significant public health concern. As the predominant mosquito species responsible for the transmission of these viruses in nearly all dengue-endemic countries, it plays a critical role in global vector-borne disease research. Unlike malaria vectors, which include a complex of species, *Ae. aegypti* exists as a single species worldwide. Only two subspecies have been identified based on morphological analyses in Africa namely, *Ae. aegypti formosus* and *Ae. aegypti aegypti* (Abuelmaali et al., 2022; Moore, 1979). *Ae. aegypti formosus* is confined to sub-Saharan Africa, while *Ae. aegypti aegypti* (hereafter called *Ae. aegypti*) is the one found globally.

The genetic diversity within *Ae. aegypti* populations is highly extensive, which has significant implications for its vectorial capacity and responses to insecticides. Global microsatellite analysis has revealed two major lineages of *Ae. aegypti* - one originating in Africa and the other outside of Africa (Gloria-Soria et al., 2016). Even within non-African populations, there is substantial genetic diversity observed across countries. Similarly, mitochondrial gene analysis has identified two distinct lineages associated with West and East Africa, respectively. These lineages have dispersed globally and now coexist in populations worldwide. Genetic variations among mosquito populations have shown significant impact on differences in transmission efficiency and species’ susceptibility to insecticides as well as the potential for adaptive traits in different environments (Bonica et al., 2018; Souza-Netoa et al., 2019; Thornton et al., 2020; Vega-Rúa et al., 2014).

To characterize the whole mitochondrial genome diversity of *Ae. aegypti*, we sequenced the complete mitochondrial genomes of nine samples from India and analyzed them alongside 34 genomes from various countries available in GenBank. Our analysis identified two distinct clusters among the Indian samples, separated by approximately 127 SNPs, with over 90% of these located in protein-coding genes (PCGs). Further investigation revealed an association between one of the clusters and *Wolbachia*, a maternally inherited endosymbiotic bacterium. While the association between mitochondrial variations and *Wolbachia* is not a new phenomenon, given the maternal inheritance of both entities, it has important implications for *Wolbachia-*based *Ae. aegypti* control strategies.

## Materials and Methods

### Mitochondrial genome sequencing from *Ae. aegypti* samples

Mitochondrial genome sequences for this study were retrieved from whole genome resequencing data from our recent study (Gupta et al., 2024). Nine samples including four normal type and five variant types (Table S1) based on morphological identification as explained in Kumar et al., 2022 were used for this analysis. Low-coverage genome sequencing of these samples was performed. The procedure for sequencing is detailed in our previous study (Gupta et al., 2024). Briefly, the raw reads obtained after illumine sequencing were trimmed for removing adapters and filtering low-quality reads using Trimgalore v0.4.04 (Krueger 2015). The adapter-clipped, high-quality reads were used for mitochondrial genome assembly using NOVOPlasty (Dierckxsens et al. 2017) in Galaxy (usegalaxy.eu)(Giardine et al., 2005), and mitochondrial genome of *Ae. aegypti* reference strain (NC_035159.1) was used as the reference. Mitochondrial genes were annotated using MITOS (Bernt et al. 2013). The circular map of the mitogenomes was visualized using Proksee (Grant et al. 2023).

### Mitochondrial genome analysis: variants identification and phylogenetics

Mitochondrial genome sequences from *Ae. aegypti* from GenBank were analysed along with our samples. The sequences were aligned in MEGA (Kumar et al., 2008) using MUSCLE (Edgar, 2004) and variants were identified. Nucleotide sequences for all the protein coding genes were extracted using reference genome sequence as a base. For phylogenetic analysis, the best substitution model was determined using Model Test in MEGA, and a maximum likelihood tree was generated with 500 bootstrap replications.

Additionally, sequences for four mitochondrial genes COI, COII, ND5, and ND4 were retrieved from GenBank and analyzed as described above. All the sequences with Ns and the ones retrieved from the mitochondrial genome sequences were removed. To obtain the maximum length of COI gene, a ∼1000 bp region of COI was selected that contained 58 sequences. COII and ND5 had ∼570bp and ∼350bp length, respectively for 54 samples each. For ND4, we retrieved 472 sequences for approximately 220 bp. To improve the visualization of the ND4 phylogenetic tree, only haplotypes were used for constructing the tree. The haplotypes were generated using DnaSP (Librado and Rozas, 2009). The phylogenetic tree for all the mitochondrial genes were constructed using Neighbor-joining method incorporated in MEGA with 500 bootstraps.

## Results & discussion

### Mitochondrial genome variations among Indian *Ae. aegypti* samples

The mitochondrial genome assemblies of nine *Ae. aegypti* samples from India ranged from 15,730 bp (M6) to 16,374 bp (M8), with an average length of 16,058 bp (Table S1; mitochondrial sequences submitted to Genbank). Five of these samples exhibited circular mitochondrial genomes (Table S1). The circular genome of one sample (M8) is shown in fig 1. The average GC content across the genomes was 21.4%, consistent with values commonly observed in insect mitochondrial genomes (Andreason et al., 2023; Do Nascimento et al., 2021; Kyriacou et al., 2024). All nine mitogenomes contained 37 genes, including 22 genes encoding tRNAs, two genes for rRNAs, and 13 PCGs. Gene lengths were generally consistent with the reference genome (NC_035159.1), except for a 138 bp deletion in all the samples, between positions 14526 bp and 14663 bp which is the part of 12S rRNA. This deletion has also been reported in samples from California and South Africa (Schmidt et al., 2018).

**Figure 1.**
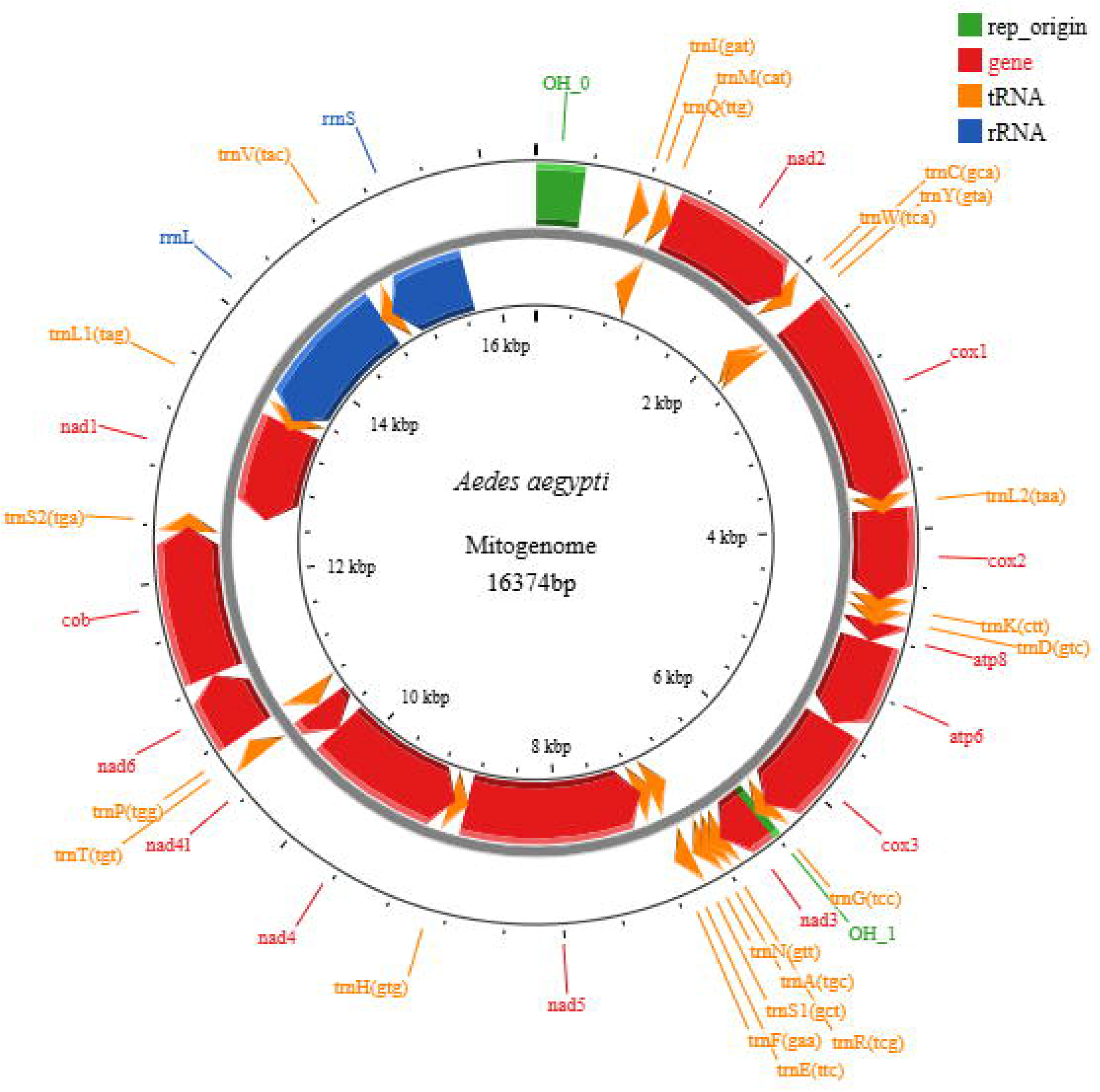
Circular map of the *Aedes aegypti* mitochondrial genome. The gene are color-coded. Genes drawn in the inner circle are transcribed anti-clockwise and the ones on the outer circle are transcribed clock-wise.

A total of 199 variants were identified in ∼14,863 bp length of mitogenome that was common across all the nine samples. This region included all the 37 genes and a portion of control region. Out of 199 variants, 197 were single nucleotide polymorphisms (SNPs) and two indels. Out of total 197 SNPs, 170 were parsimony informative and 3 were heteroplasmic. Majority of the SNPs (90%) were found in the PCGs (Table 1). Eight SNPs and one indel were detected in tRNA genes. The 12S rRNA gene exhibited four SNPs and one indel, while the 16S rRNA gene showed seven SNPs and one indel (2bp). One variation was found in the intergenic region between tRNA gene and ND3. The variations, particularly in the tRNA and rRNA genes are generally rare and could potentially impact mitochondrial translation and ribosomal function, both of which are critical for mitochondrial activity and overall mosquito fitness (Dannfald et al., 2021).

**Table 1.**
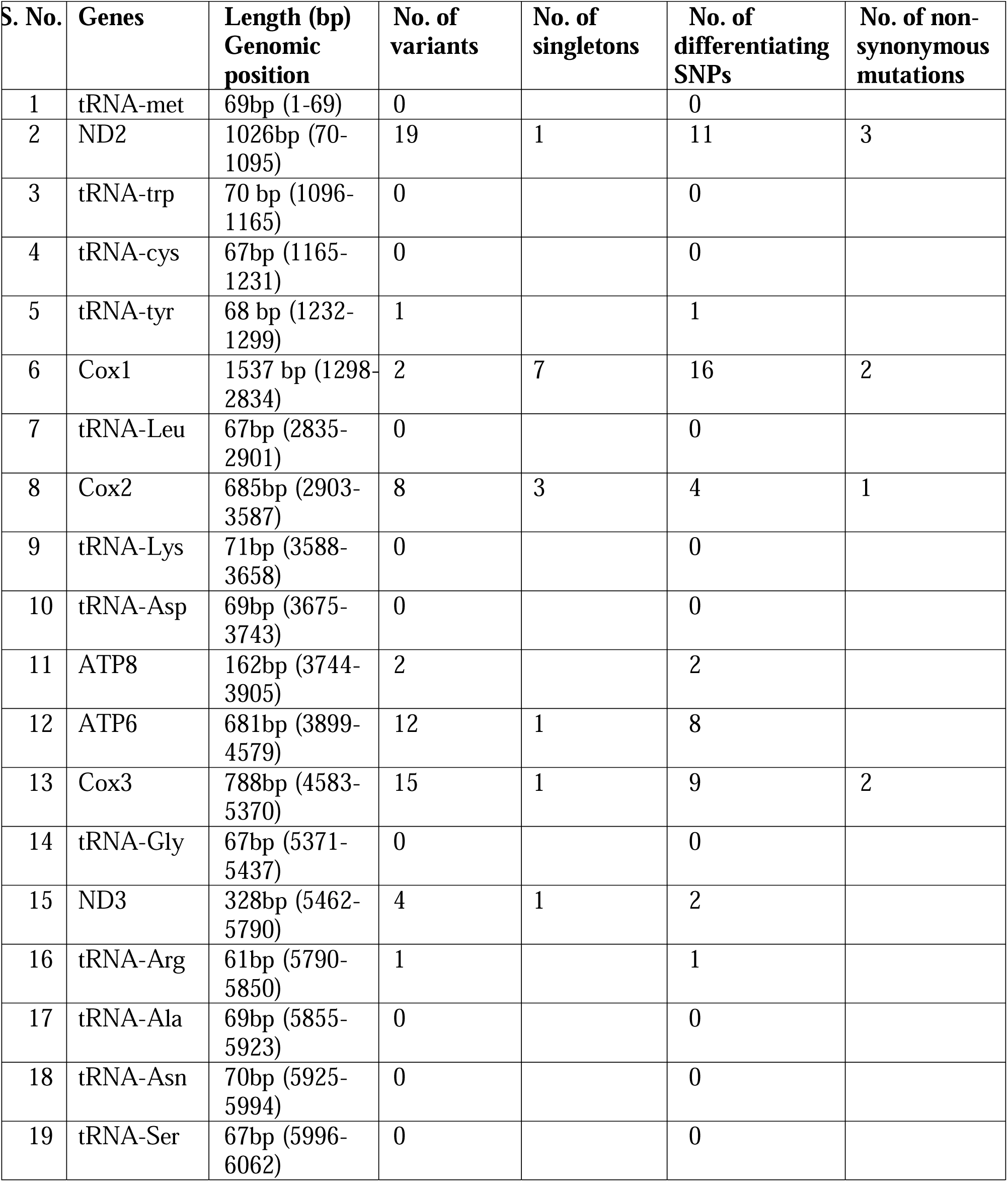
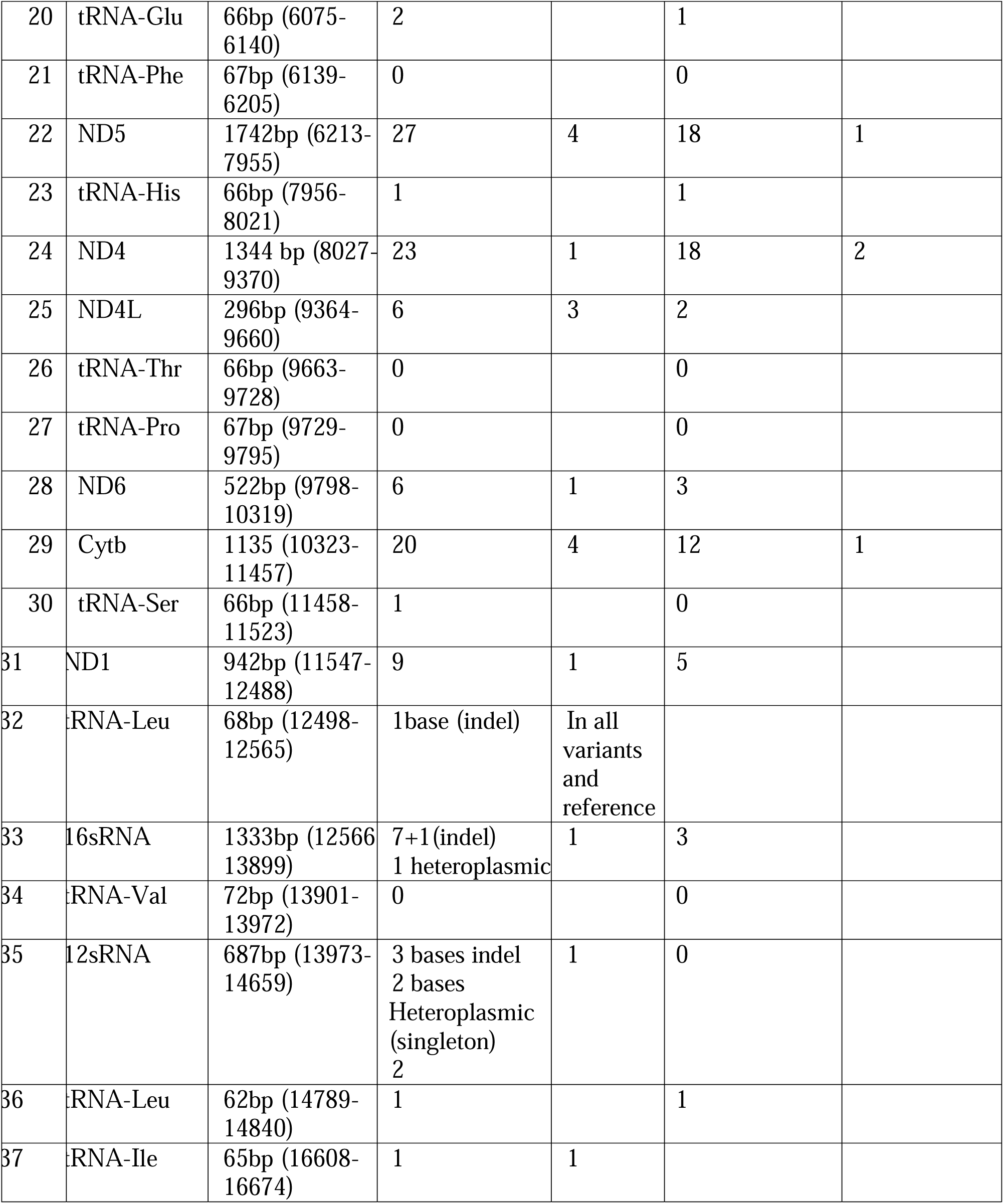
Mitochondrial genes, their length, and the number of variations, singletons, and non-synonymous mutations found in nine mitochondrial genomes of *Aedes aegypti* samples from India.

Among the PCGs, the CO1 gene exhibited the highest single nucleotide polymorphism (SNP) count (29), followed by ND5 (27) and ND4 (23) (Table 1). Majority of these SNPs were neutral as only 11 were classified as non-synonymous. However, mitochondrial genes tend to be highly conserved due to their critical roles; even minor mutations can result in significant adverse effects (Anderson et al., 2022; Ballard et al., 2007; Camus et al., 2017; Cañadas-Garre et al., 2024). Mutations in these genes can impact the electron transport chain, leading to changed ATP production or increased oxidative stress (Bhatti et al., 2017; Guo et al., 2013; Kowalczyk et al., 2021). This oxidative stress can, in turn, influence various aspects of mosquito biology and behavior, including metabolism, reproductive success, and overall fitness. Although most variations in mitochondrial genomes are neutral, they can become fixed within populations and form specific haplogroups. Sometimes, these haplogroups have shown strong associations with various traits or phenotypes (Hendrickson et al., 2008; Kenney et al., 2013; Kuroki and Fukami, 2023; Mayer et al., 2020; Shaikevich et al., 2016). However, the biological significance of variations in cellular processes and overall mosquito growth of *Ae. aegypti*, functional characterisation of these variations remains to be determined.

### Phylogenetic clustering of *Ae. aegypti* mitogenomes

Phylogenetic analysis of the nine Indian *Ae. aegypti* samples and reference genomes revealed two distinct clades (Fig. 2). Clade I included six samples, along with the reference genome, while Clade II included three samples. There was no clear genetic separation between the two *Ae. Aegypti* morphotypes as observed in nuclear genome analysis (Gupta et al., 2024), likely due to interbreeding between the two types. Notably, more than half of the total SNPs (127 of the 197) distinguished the samples from the two clades. Among these, 112 were found in PCGs indicating a significant number of variations that could potentially alter the genetic characteristics and biology of this vector species. Sample M8 shared variants with both clades and was more closely aligned with Clade I. This could potentially be due to recombination or the emergence of a new lineage driven by local environmental factors.

**Figure 2.**
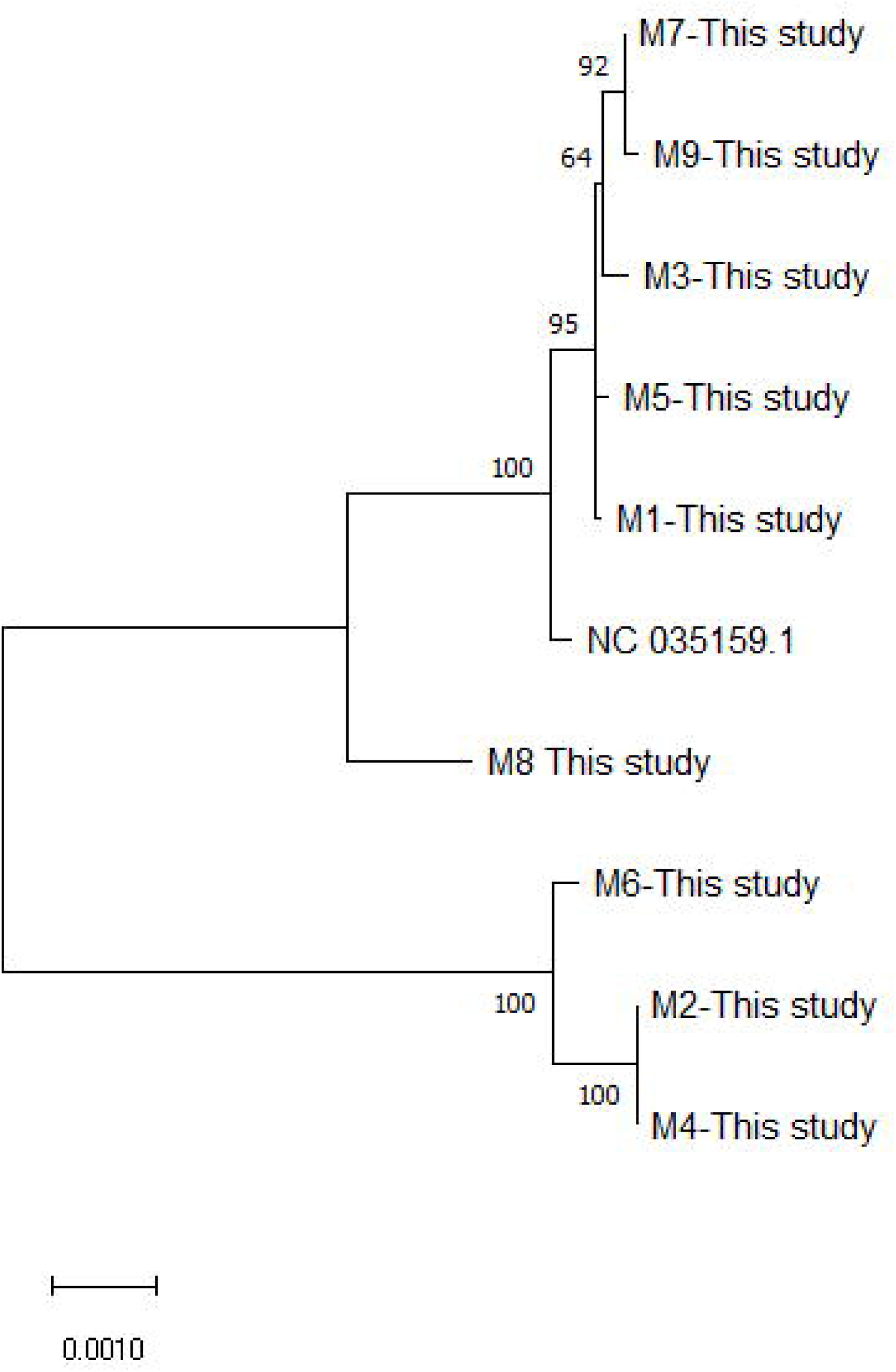
Phylogenetic tree of the mitochondrial genomes from nine *Aedes aegypti* samples from India, constructed using the Maximum Likelihood method with 500 bootstrap replicates.

To place these patterns in a global context, we compared our mitochondrial genome data with samples from other countries available in public databases. Phylogenetic analysis of 43 genomes identified two clades (Figure 3). The largest clade consisted of 38 genomes from diverse global regions, including the reference genome, reflecting a widespread lineage. The second clade was specific to five samples from Saudi Arabia. Notably, the largest clade was further divided into two sub-clades. These two clades aligned with the clustering of our Indian samples as shown in Fig 2. One of these sub-clades (clade I) contained samples from India, the USA, Brazil, Saudi Arabia, including the reference genome and the Rockefeller strain, while the other clade (clade II) included three Indian samples alongside those from Australia, Saudi Arabia, and South Africa (Figure 3). This clustering of global samples into two major clades aligns with earlier findings from various studies that identified two primary lineages based on mitochondrial markers, indicating their global spread and adaptation (Abuelmaali et al., 2021; Lima and Scarpassa, 2009; Maitra et al., 2019; Soghigian et al., 2020).

**Figure 3.**
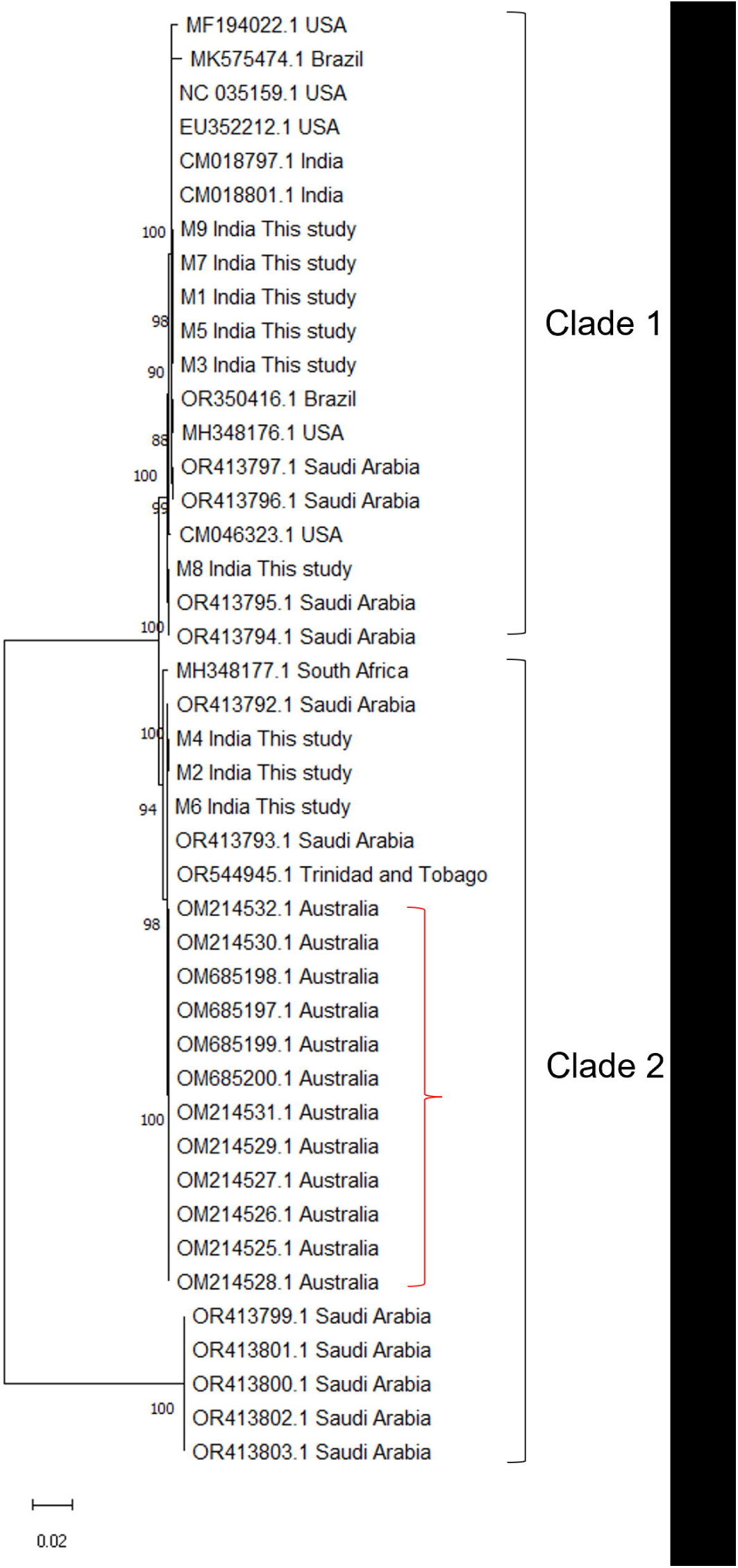
Phylogenetic tree of the mitochondrial genomes from global *Aedes aegypti* samples, constructed using the Maximum Likelihood method with 500 bootstrap replicates. The samples shown within red bracket are *Wolbachia* infected.

Clustering analysis revealed similar patterns across four traditional markers; COI, COII, ND4 and ND5, identifying two major clades consistent with the clustering observed in mitogenome samples (Fig S1-S4). The consistent clustering pattern across these markers highlights the reliability of these traditional markers in tracing genetic variations and the evolutionary history of this mosquito species and indicates that the mitochondrial genes of *Ae. aegypti* share a common genealogy.

The existence of well-established global lineages highlights the potential biological significance of each clade, as they have persisted in these areas long after their initial invasion. Variations in specific mitochondrial genes have been associated with various diseases in humans (Cañadas-Garre et al., 2024; Gusic and Prokisch, 2021; Wei and Chinnery, 2020). Additionally, distinct mitochondrial lineages have been correlated with climatic factors and geographical locations, suggesting that environmental pressures may influence mitochondrial evolution (Balloux et al., 2009; Maca-Meyer et al., 2001; Zhao et al., 2024). Given that mitochondrial genes play a critical role in regulating temperature and energy metabolism, these associations are plausible. Understanding how these genetic variations interact with environmental factors can provide valuable insights into the adaptive strategies of these lineages and their implications for ecology and evolution.

### Association of mitochondrial variations with *Wolbachia* infection

We made an intriguing observation during our global mitogenome analysis that the *Ae. aegypti* samples infected with *Wolbachia* were exclusively found within clade II (Figure 3). However, the status of infection of all the samples is not known, the infected samples were confined to one clade only. This finding is similar to the analysis of a recent study by Thia and coworkers where all infected samples were found in one clade, except for a single *Wolbachia*-infected sample from Malaysia, which was separated from the infected samples from Australia (Thia et al., 2024). Since genome sequences used in this study from Ross et al. (2021) were unavailable, we could not include them in the analysis (Ross et al., 2021). However, the variant haplotypes available in the 750 bp fragment of COI from the same study indicated that the *Wolbachia*-infected Malaysian samples aligned with Clade II, while the uninfected sample from Malaysia had a different haplotype aligning with clade I.

To explore this further, we analyzed sequences of COI, ND4, ND5, and other three fragments published by Yeap et al. (2016), which included samples from various countries, including *Wolbachia*-infected samples from Australia (Yeap et al., 2016). These sequences also displayed the two clades, with all infected samples grouped within one clade, following the pattern observed in our full mitogenome analysis (Fig S5-S10). However, it should be noted that these samples were artificially infected with the wMel or wAlb strain of *Wolbachia*. It could be argued that the original mosquito female lines used for transfection contained these mitochondrial lineages, which were maintained through subsequent generations. This is plausible, as tight maternal inheritance of *Wolbachia* has been documented by Huang and co-workers who identified only three haplotypes differing by a single nucleotide position eight years post-release of *Wolbachia*-infected mosquitoes in Australia (Huang et al., 2020).

To support this, we searched literature of the mitochondrial sequences front eh mosquito samples found *Wolbachia*-positive from field populations. We retrieved COI sequences from two such studies published from Taiwan (Chao and Shih, 2023) and Manila (Reyes et al., 2024), and analysed them along with our samples. These sequences exhibited a similar pattern, showing that *Wolbachia*- infected samples clustered within Clade II (Figure 4). However, this cade also contained uninfected samples (Chao and Shih, 2023) which can be explained by two possible scenarios. One possibility is that both the lineages were originally infected but one of them subsequently lost the bacteria due to imperfect transmission as observed in *Drosophila melanogaster* where 1-3% of the progenies of *Wolbachia*-infected lines were found uninfected (Carrington et al., 2011; Hague et al., 2020).

**Figure 4.**
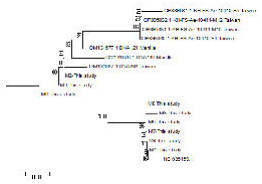
Neighbor joining tree for COI sequences from field samples of *Ae. aegypti* infected with *Wolbachia* collected from Taiwan and Manilla, Philippines along with Indian samples. The numbers shown on the branches are bootstrap values.

Alternatively, *Wolbachia* may have been recently introduced into *Ae. aegypti* and possibly had a reproductive advantage within only one lineage. This particular mitochondrial background might be more conducive to *Wolbachia* colonization or replication, which could explain the lower occurrence of this symbiont in natural *Ae. aegypti* populations. wAlbB transfection was found successful in the mosquitoes having Australian mitochondrial haplotype (Ross et al., 2021), demonstrating the potential for effective *Wolbachia* colonization when paired with a suitable mitochondrial background.

*Wolbachia* is the most commonly found endosymbiont in mosquitoes. Although it was once widely believed that *Ae. aegypti* did not harbor *Wolbachia* (Kittayapong et al., 2000), several reports on *Wolbachia* infection in the natural *Ae. aegypti* populations have been published in the last decade (Balaji et al., 2019; Bennett et al., 2019; Carvajal et al., 2019; Coon et al., 2016; Kulkarni et al., 2019; Teo et al., 2017; Thongsripong et al., 2018; Vinayagam et al., 2023; Zhang et al., 2022).

Notably, a large-scale screening of 2663 samples across 27 populations from multiple continents did not detect any *Wolbachia* infections (Gloria-Soria et al., 2018). This raises the possibility that *Wolbachi*a infected samples identified in these countries have either originated from laboratory strains or lacked confirmed *Wolbachia* infection, as suggested by Ross and Hoffman (Ross and Hoffmann, 2024). The ‘Eliminate Dengue project’, which involves the release of laboratory reared *Wolbachia*-infected *Ae. aegypti* started field trials in Australia and then extended to 11 other countries since 2011 (Hoffmann et al., 2011). The initial reports of *Wolbachia* infections in wild *Ae*.

*aegypti* populations started emerging after the release (Balaji et al., 2019; Baldini et al., 2014; Bennett et al., 2019; Carvajal et al., 2019; Hegde et al., 2018; Kulkarni et al., 2019; Teo et al., 2017; Thongsripong et al., 2018). The global spread of *Aedes* mosquitoes from experimental areas or countries through travel infrastructure is a real possibility (Eritja et al., 2017; Reiter and Sprenger, 1987). However, to definitively rule out the invasion of Australian strains via travel, whole-genome analyses comparing infected samples with Australian genomes would be essential. Such a comparison would help identify any shared genetic markers or haplotypes that could indicate recent migration or introduction of these strains into these countries. In our nine samples, we could not identify reads matching the *Wolbachia* genome, which may be attributed to low coverage sequencing data or the absence of infection in our samples. Notably, *Wolbachia* has been detected in natural populations of *Ae. aegypti* in India as early as 2019 (Balaji et al., 2019; Vinayagam et al., 2023), prior to the initiation of experiments involving *Wolbachia*-infected mosquitoes in the country (Gunasekaran et al., 2022; Sadanandane et al., 2022).

Hitchhiking of mitochondrial lineages with maternally inherited symbionts like *Wolbachia* has been observed in many other organisms (Bakovic et al., 2020; Charlat et al., 2009; Jiang et al., 2018, 2014; Kirik et al., 2020; Shastry et al., 2022; Sucháčková Bartoňová et al., 2021). For example, a study conducted by Jiang et al, identified three major clades of *Polytremis nascens* based on mitochondrial haplotypes; one of the clades comprising *Wolbachia* infected butterflies and the other two with uninfected butterflies (Jiang et al., 2014). Similarly, Rasgon et al, identified two clusters of mitogenomic variation in *Culex pipiens*, one contained uninfected haplotypes and the others were infected (Rasgon et al., 2006). Although the data on these clusters has not been analyzed for associations, a significant number of variations in the ND4 fragment has been documented.

Moreover, the presence of *Wolbachia* is reported to heighten the prevalence of cryptic species within the *Culex pipiens* complex (Dumas et al., 2016). It is possible that these variations may not be supporting the propagation of *Wolbachia* within mosquito populations. This raises intriguing questions about the role of *Wolbachia* in shaping mitochondrial diversity and functionality. It also opens avenues for exploring how these symbiotic relationships influence the evolutionary trajectory of the host’s mitochondrial genes.

While there could be other unknown factors or symbionts influencing these patterns, it is possible that specific mitochondrial variations are adaptations to *Wolbachia* presence. Previous studies have shown that *Wolbachia* can induce oxidative stress in host mosquitoes (Pan et al., 2018, 2012). This stress could negatively impact the host’s fitness; however, it is likely that the mitochondria undergo adaptations that enable them to thrive. Future research is warranted to explore these interactions to further elucidate their roles in vector biology and its implications in vector control through *Wolbachia* infection.

In conclusion, this study provides a comprehensive and global perspective on the mitochondrial genetic variation in *Ae. aegypti*. The data offers a reference base for future research aimed at understanding the evolutionary dynamics and functional significance of mitochondrial diversity in mosquito populations. Our analysis identified two lineages based on complete mitochondrial genome which were previously reported using traditional mitochondrial markers, highlighting that these markers are good representations of the mitochondrial genome diversity and the evolutionary history.

Additionally, this study highlights a potential link between mitochondrial genetic background and *Wolbachia* infection, opening new avenues for research into host-symbiont interactions. Future research should focus on the following directions: i) investigating whether *Wolbachia* can thrive in *Ae. aegypti* strains with the other mitochondrial lineage ii) determining if these two strains are reproductively isolated due to *Wolbachia* infection iii) exploring functional implications of mitochondrial variations on mosquito biology and their interactions with *Wolbachia*, and iv) understanding how the existence of these two strains globally might affect *Wolbachia*-based mosquito control efforts. Understanding these dynamics could provide valuable insights for *Wolbachia*-based vector control strategies. If *Wolbachia* can survive only in the mosquitoes with one type of lineage, the presence of the mosquitoes with other lineage in that geographical area would compromise the impact of *Wolbachia*-based control efforts.

## CRediT authorship contribution statement

**Bhavna Gupta:** Conceptualization, Formal analysis, Funding acquisition, Methodology, Project administration, Supervision, Visualization, Writing – original draft. **Melveettil Kishor Sumitha:** Data curation, Writing – review and editing. **G Navaneetha Pandiyan:** Data curation, Writing –

review and editing. **Mariapillai Kalimuthu:** Data curation, Writing – review and editing. **Rajaiah Paramasivan:** Resources. **Manju Rahi:** Resources.

## Ethics statement

Ethical clearance was not required for this work. However, informed consent was obtained from the owners of the households where mosquito collections were conducted. All procedures were carried out with respect for the local community and environmental considerations.

## Acknowledgements

The authors would like to thank Indian Council of Medical Research (ICMR), New Delhi for the intramural support. Melveettil Kishor Sumitha and G Navaneetha Pandiyan would like to thank Madurai Kamaraj University for supporting their research.

## Supplementary figure legends

**Supplementary** Figure 1: Phylogenetic tree of COI sequences of our study along with the sequences retrieved from GenBank, constructed using the Maximum Likelihood method with 500 bootstrap replicates.

**Supplementary** Figure 2: Phylogenetic tree of COII sequences of our study along with the sequences retrieved from GenBank, constructed using the Maximum Likelihood method with 500 bootstrap replicates.

**Supplementary** Figure 3: Phylogenetic tree of ND4 sequences (haplotypes were used) of our study along with the sequences retrieved from GenBank, constructed using the Maximum Likelihood method with 500 bootstrap replicates.

**Supplementary** Figure 4: Phylogenetic tree of ND5 sequences of our study along with the sequences retrieved from GenBank, constructed using the Maximum Likelihood method with 500 bootstrap replicates.

**Supplementary** Figure 5: Phylogenetic tree of COI sequences of our study along with the sequences published by (Yeap et al., 2016) constructed using the Maximum Likelihood method with 500 bootstrap replicates.

**Supplementary** Figure 6: Phylogenetic tree of ND4 sequences of our study along with the sequences published by (Yeap et al., 2016) constructed using the Maximum Likelihood method with 500 bootstrap replicates.

**Supplementary** Figure 7: Phylogenetic tree of ND5 sequences of our study along with the sequences published by (Yeap et al., 2016) constructed using the Maximum Likelihood method with 500 bootstrap replicates.

**Supplementary** Figure 8: Phylogenetic tree of the sequences of our study along with the sequences amplified by Primer 1 published by (Yeap et al., 2016) constructed using the Maximum Likelihood method with 500 bootstrap replicates.

**Supplementary** Figure 9: Phylogenetic tree of the sequences of our study along with the sequences amplified by Primer 4 published by (Yeap et al., 2016) constructed using the Maximum Likelihood method with 500 bootstrap replicates.

**Supplementary** Figure 10: Phylogenetic tree of the sequences of our study along with the sequences amplified by Primer 6 published by (Yeap et al., 2016) constructed using the Maximum Likelihood method with 500 bootstrap replicates.

