## Supplementary figures and images for "Whole mitochondrial genome analysis of *Aedes aegypti* reveal association with *Wolbachia* infection"

### Supplementary figure 1

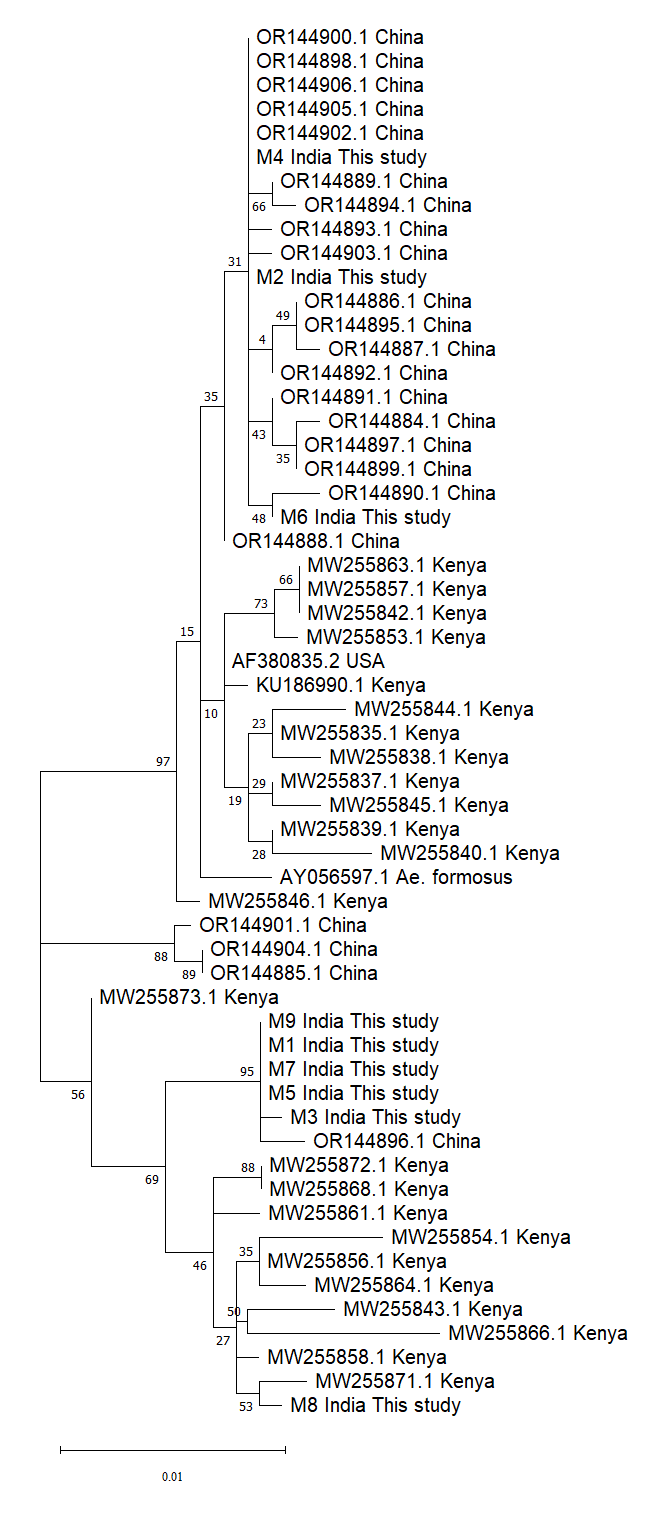

### Supplementary figure 2

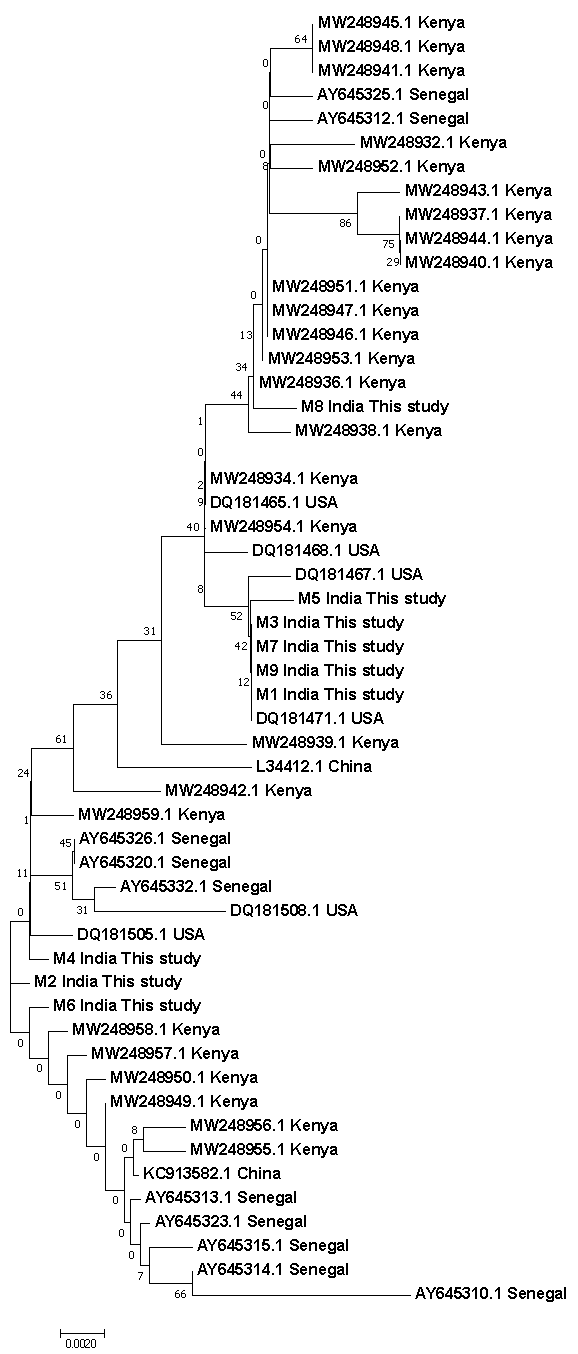

### Supplementary figure 3

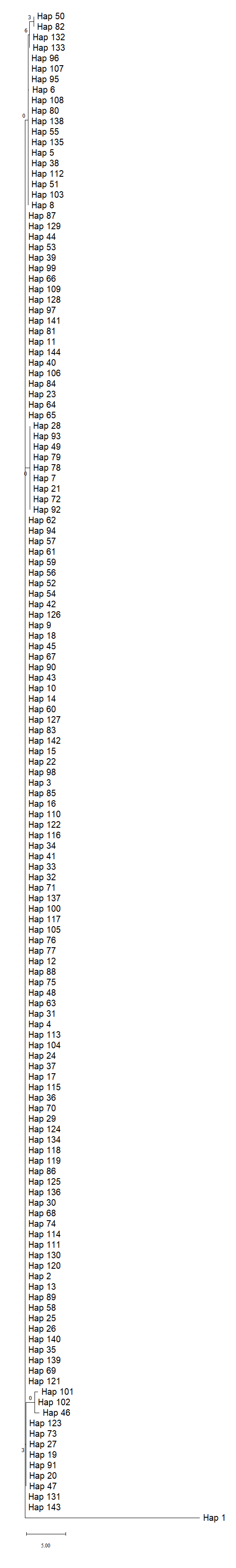

### Supplementary figure 4

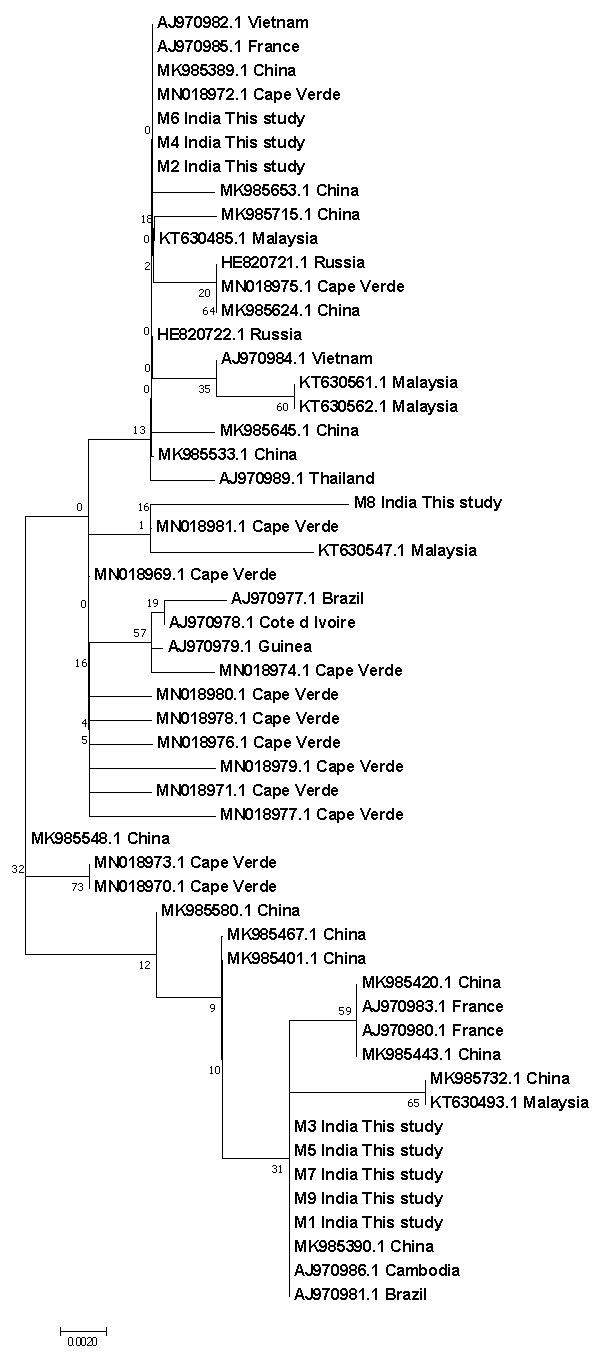

### Supplementary figure 5

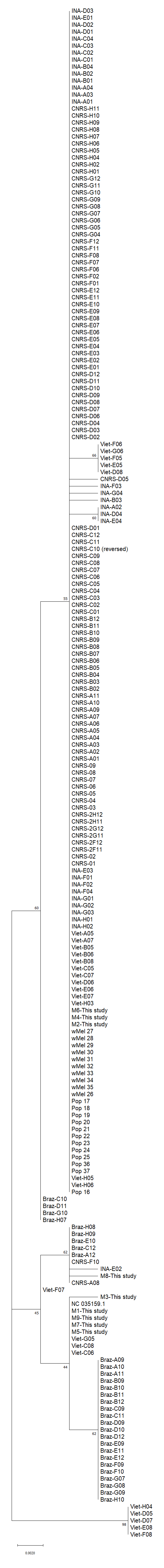

### Supplementary figure 6

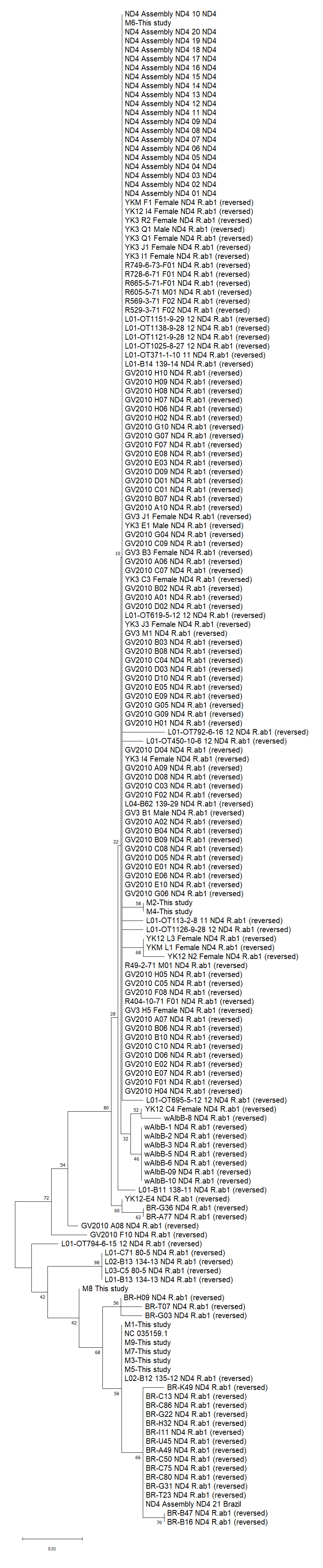

### Supplementary figure 7

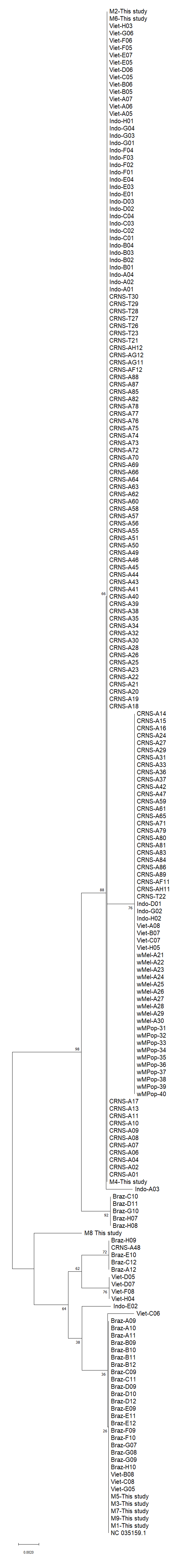

### Supplementary figure 8

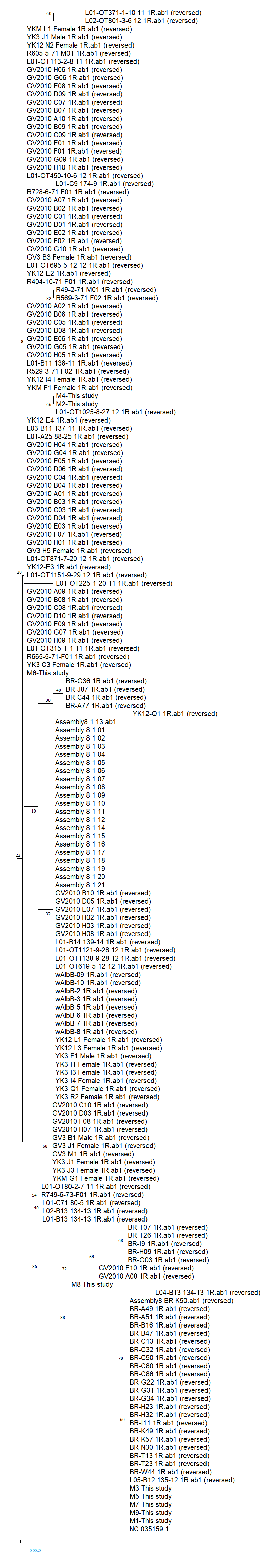

### Supplementary figure 9

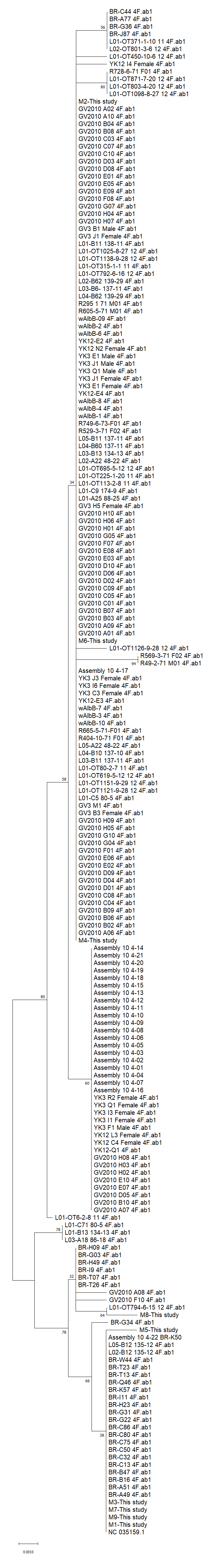

### Supplementary figure 10

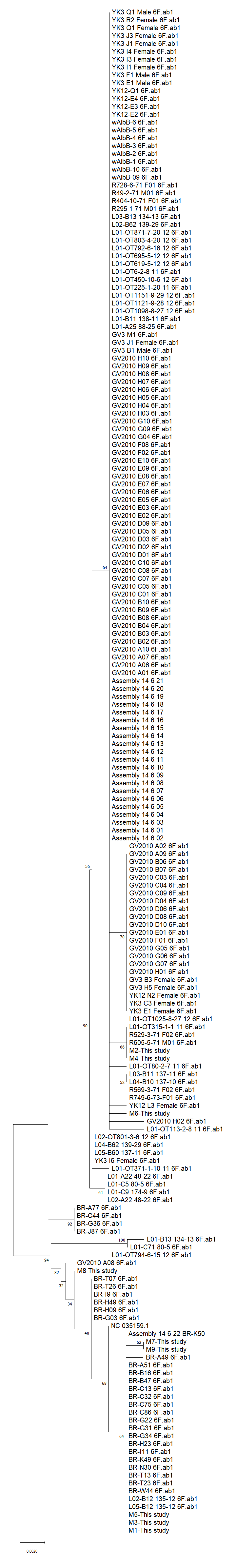
