## Supplementary table for "Whole mitochondrial genome analysis of *Aedes aegypti* reveal association with *Wolbachia* infection"

**Table S1.** Details of mitochondrial genome assembly for the seven *Aedes aegypti* samples from India.

| **Sample No. (Morphotype)** | **Mitogenome size (bp)** | **Circularity** | **GC content (%)** |
| --- | --- | --- | --- |
| M1 (Variant) | 16010 | No | 21.6 |
| M2 (Normal) | 15973 | No | 21.56 |
| M3 (Normal) | 16117 | Yes | 21.43 |
| M4 (Normal) | 16073 | Yes | 21.45 |
| M5 (Normal) | 16081 | No | 21.47 |
| M6 (Variant) | 15730 | Yes | 21.84 |
| M7 (Variant) | 16081 | No | 21.47 |
| M8 (Variant) | 16374 | Yes | 21.09 |
| M9 (Variant) | 16081 | No | 21.47 |
